# Molecular epidemiology of the globally spreading genetic lineage IV of peste des petits ruminants virus

**DOI:** 10.64898/2026.05.18.725933

**Authors:** Maxime Courcelle, Kadidia Tounkara, Samuel Mantip, Mamadou Niang, Cheick Abou Kounta Sidibe, Amadou Sery, Marthin Dakouo, Pam Dachung Luka, Adeyinka Adedeji, David Shamaki, Maryam Muhammad, Yahia Hassan Ali, Intisar Kamil Saeed, Joseph Awuni, Theophilus Odoom, Patrick Ababio Tetteh, Daniel Tawiah Yingar, Abel Wade, Simon Dickmu, Aissatou Diddi, Hazar Shawash, Emmanuel Couacy-Hymann, Koffi Yao Mathurin, Hatem Ouled Ahmed Ben Ali, Sonia Ben Hassen, Salma Haddouchi, Ahmad Alm-ajali, Tirumala B.K. Settypalli, Charles E. Lamien, Habib Salami, S. Rassoul, Mohadji Asnaoui, Catherine Cetre-Sossah, Samia Guendouz, Olivier Kwiatek, Geneviève Libeau, William G Dundon, Arnaud Bataille

## Abstract

Peste des petits ruminants (PPR) is a highly contagious viral disease of small ruminants caused by the peste des petits ruminants virus (PPRV), which is classified into four distinct genetic lineages (I–IV). A critical concern in the recent epidemiological history of PPRV is the rapid and widespread expansion of lineage IV (LIV) across West Africa over the past decade. This dominance suggests a potential adaptive advantage of circulating LIV strains in the region’s current epidemiological context. In this study, we obtain the genome sequence of 26 new PPRV samples, including historical (pre-2000) and many recent African LIV isolates, offering the first opportunity to investigate the evolutionary history of LIV in Africa and identify genetic events potentially associated with its recent spread. Phylogenomic analyses implemented on a dataset of 167 curated PPRV genome sequences reveal that the most ancestral LIV group comprises strains circulating in Sub-Saharan Africa (designated clade LIVssa), providing robust evidence for an African origin of lineage IV. Our results further indicate that PPRV strains linked to the recent West African expansion of LIV belong to a specific LIVssa subgroup, termed NigB. We identified multiple signatures of selection pressure within the LIVssa sublineage, particularly in the NigB cluster. Several amino acid substitutions unique to LIVssa or NigB were detected, some of which may impact protein function and warrant prioritised investigation. Additional genomic data are required to confirm the association between the NigB group and the ongoing spread of LIV in West Africa. The evolutionary adaptations observed in LIVssa - potentially enhancing transmission efficiency, host range or pathogenicity - could undermine current disease control strategies in regions where PPR poses significant threats to food security and local economies.

**Author Summary:** Peste des petits ruminants virus (PPRV) infects sheep and goats across Africa, Middle East, Asia and Europe, causing disease with major impact on global economy and food security. One genetic lineage of PPRV, called lineage IV (LIV), is at the origin of most recent expansion of the distribution of the disease, including replacement of other lineages in areas of African where PPRV is historically present. Here, we generated genome sequences from PPRV LIV isolates from different dates and places to study the evolution of this genetic lineage and explore whether its recent spread can be associated with the appearance of new mutations in the virus genome. Our results provide evidence that the PPRV LIV originated in Sub-Saharan Africa and identify mutations present only virus isolates currently spready in new regions of Africa. Further research should investigate the impact of these mutations on protein functions and capacity of transmission of PPRV.

## Introduction

Peste des petits ruminants is a highly contagious disease of small ruminants that can rapidly spread across national borders, causing significant economic consequences. It is considered a global priority among Transboundary Animal Diseases (TADs) and is currently the target of a global eradication effort led by the World Organisation for Animal Health (WOAH) and the Food and Agriculture Organization of the United Nations (FAO) (*1, 2*). The recent emergence of the disease in Europe highlights the risk of spread in PPR-free countries from endemic areas through legal and illegal animal movements (*3*) and the difficulty of controlling this disease.

The disease is caused by peste des petits ruminants virus (PPRV), a morbillivirus (family: *Paramyxoviridae*) closely related to measles and canine distemper viruses. This virus has a single-stranded negative sense RNA genome of close to 16,000 nucleotides, coding for 6 structural proteins and 2 non-structural proteins. Similar to other RNA viruses, PPRV is characterised by a fast evolutionary rate [between 10^-3^ and 10^-4^ substitutions per site per year (*4*)] due to its error-prone RNA polymerase (*5*). This high mutational capacity and global endemic distribution across Africa, the Middle East and Asia, has led to a high genetic diversity among circulating PPRV strains, which are now differentiated into 4 distinct genetic lineages (lineages I, II, III and IV). Lineage IV is the most genetically diverse and globally spread, while the other three lineages are currently restricted to sub-Saharan Africa (*4*). PPRV strains vary widely in their virulence across different species and breeds (*6, 7*), but there is only one serotype and existing vaccines provide life-long protection against all circulating strains (*8*). Recent phylogenomic studies have shown that PPRV evolution in different epidemiological contexts has resulted in lineage-specific positive selection of mutations of potentially functional importance (*4*), notably in association with adaptation to atypical hosts, such as wild artiodactyls (*9*), although much remains to be explored on this question (*10*).

A major point of concern regarding the recent epidemiological history of PPRV is the widely reported spread of lineage IV (LIV) in West Africa over the past decade, replacing strains of the other three lineages where they have been circulating (*11–17*). Such a take over suggests an adaptive advantage of LIV strains circulating in West Africa in the current epidemiological context. For a long time, LIV was considered to have originated in Asia, as it was first detected in India in 1987 (*18*). However, the latest phylogenomic analyses suggest that this genetic lineage originated in sub-Saharan Africa, though available LIV genome sequences from Africa are very limited (*4, 9*). Until now, no phylogenomic study has focused on exploring the history of the emergence and evolution of LIV in sub-Saharan Africa and the possible appearance of new advantageous mutations in strains circulating in the region over the past decade. Functional changes in viral proteins associated with modification in replication capacity or host range of the virus in Africa could have an important impact on control efforts.

There is a large, long-term collaboration effort involving national reference laboratories for PPR across the world, the WOAH/FAO reference laboratories for PPR and the joint FAO/IAEA centre, notably through the WOAH network for national reference laboratories for PPR (https://www.ppr-labs-oie-network.org/). These collaborations have provided the opportunity to conduct PPRV full genome sequencing at a scale unseen before for the virus, with a special focus on sub-Saharan Africa, where data is crucially lacking (*4*). Here we present the results of genomic analyses of 26 samples of PPRV, including historical (before 2000) and recent African LIV samples, providing a first opportunity to study the evolutionary history of LIV in Africa and identify evolutionary events that could be associated with the spread of the lineage in West Africa over the past decade.

## Results

### Full genome sequencing of PPRV samples

A total of 22 complete or almost complete (>90% coverage) and 4 partial (70-89% coverage) sequences of the PPR virus genome were obtained from sheep and goat samples collected and identified as infected with PPRV by national veterinary services and other institutions in multiple countries in Africa, the Middle East and Asia within the framework of surveillance activities or previous studies (Table 1; Figure 1). Full genome sequencing and sequence data analyses were conducted at CIRAD, Montpellier, France, which serves as the WOAH/FAO and EU reference laboratory for PPR (https://eurl-ppr.cirad.fr/; https://www.ppr-labs-oie-network.org/; N=22) and at the Animal Production and Health Laboratory of the Joint FAO/IAEA Division (N=4) using their in-house protocols adapted for Illumina or Ion Torrent high-throughput sequencing (Table 1; Sequence Read Archive accession number: PRJNA717034). Sequencing efforts concentrated on genetic lineage IV PPR virus recently collected in Africa, but also included samples from the three other lineages and/or collected before 2010 (Table 1), which are still poorly represented in published full genome databases (*19*). Recombination tests were used to confirm that the new sequences obtained did not contain fragments of sequences from other strains manipulated in the laboratories (*19*), before aligning them with the annually updated, curated dataset of publicly available PPRV genome sequences, accessible via the WOAH network for PPR reference laboratories (https://www.ppr-labs-oie-network.org/). Based on this alignment, additional datasets were created for the coding region of each PPR virus gene.

**Figure 1.**
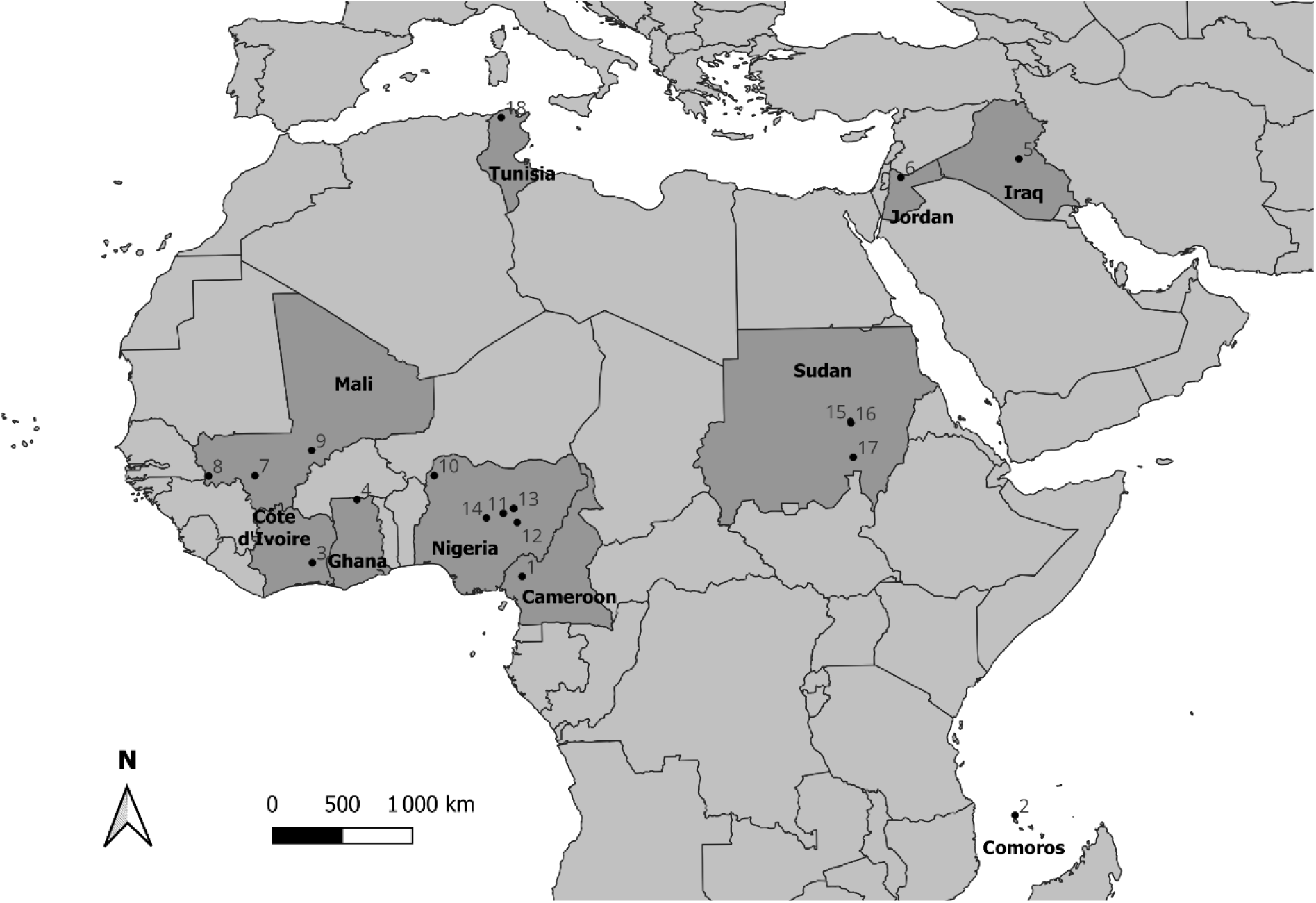
Map of the geographical location of peste des petits ruminants virus samples sequenced in this study. Numbers correspond to the name of the village or town sampled as indicated in Table 1.

**Table 1.**
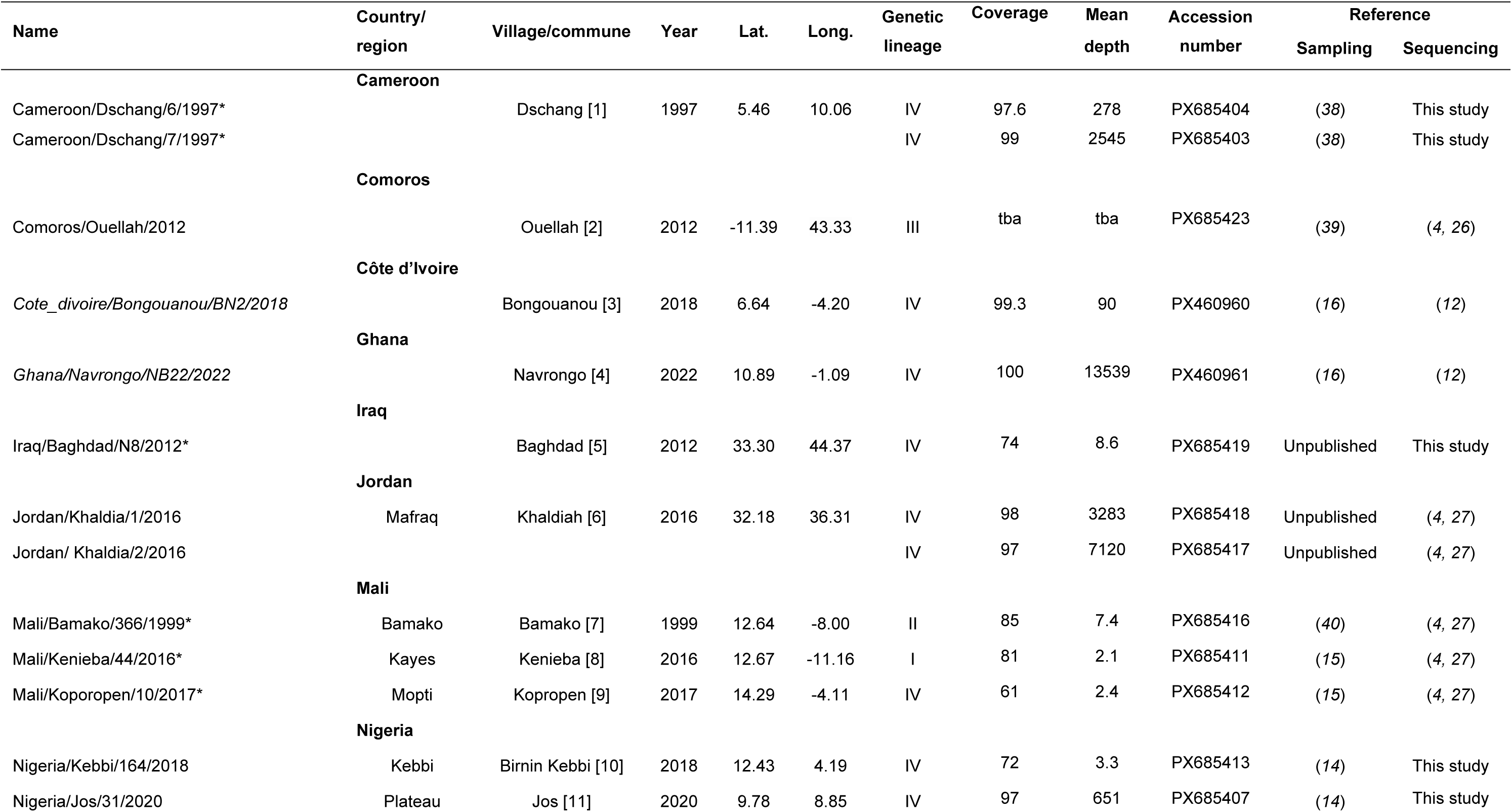
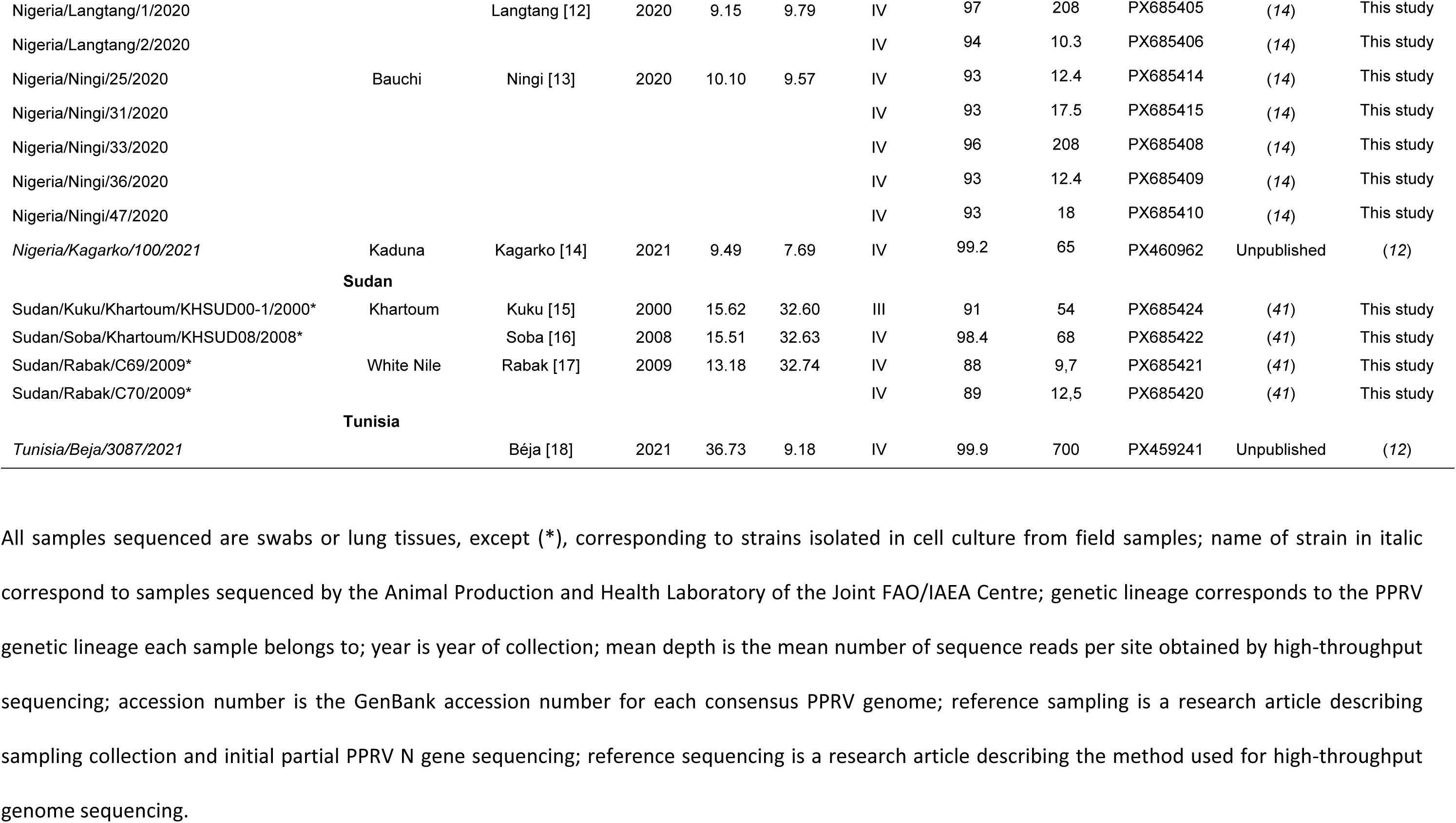
List of samples and genome sequencing results.

### Phylogenetic and phylogeographic analyses based on full genome sequences

A total of 10 genomes were obtained from samples collected in Nigeria between 2018 and 2022, a sequence from Sudan obtained in 2008 in Khartoum, and sequences from Mali (Koporopen/2017), Côte d’Ivoire (2022), and Ghana (2022) grouped with three LIV sequences already published from West and Central Africa. The sequences obtained from historical samples collected in 1997 in Cameroon placed at the root of this cluster (Figure 2). This sub-Saharan cluster (hereafter identified as LIVssa) is well separated with other geographically defined clusters within the genetic lineage IV, defined as in a previous study, as North-East Africa (including Ethiopia; LIVne), Middle East (LIVme) and Asian (LIVa) sublineages within LIV (*4*). Sequences from samples collected in the Middle East (Jordan and Iraq) clustered within LIVme as expected (Figure 2). Two sequences from Rabak in Sudan (2009) and the sequence from Tunisia (2022) clustered with the LIVne. The older sequence from Sudan (2000) grouped with older LIII sequences from Ethiopia and Sudan and the sequence from Comoros was confirmed as belonging to lineage III. The historical sequence from Mali (1999) was found to be closely related to the already published Malian sequence from that year. The sequence obtained from Kenieba in Mali (2016) was identified as belonging to lineage I, representing the most recent full genome from this PPRV lineage (Figure 2).

**Figure 2.**
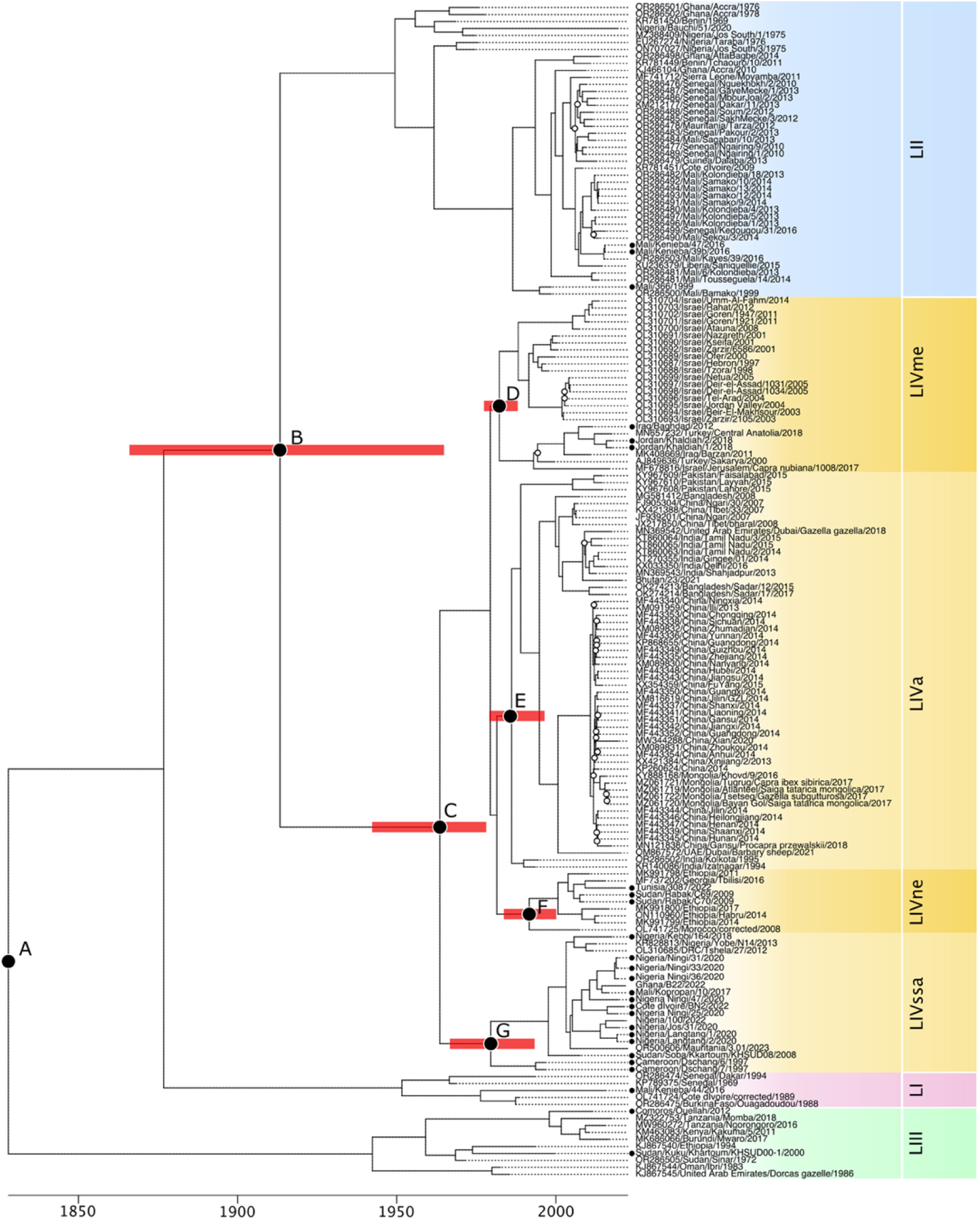
Time-calibrated Bayesian phylogenetic tree of peste des petits ruminants virus (PPRV). The x-axis represents time in years. Coloured background shading distinguishes the four PPR lineages (LI, LII, LIII, and sublineages of LIV). Filled circles indicate nodes for which the estimated age and most likely geographic origin are discussed in the main text (see Table 2); red horizontal bars denote the corresponding 95% highest posterior density intervals (HPD) for node ages. Dots indicate sequences newly generated in this study. Open circles mark nodes with posterior probability below 0.70.

**Table 2.** Estimated time to most recent common ancestor (TMRCA) for key points of the PPR phylogeny described in figure 2, with associated 95% HPD and most likely region of origin.

| Node | TMRCA | 95% HPD | Most likely origin (% probability) |
| --- | --- | --- | --- |
| (A) PPR Root | 1828 | 1744 - 1908 | Sub-Saharan Africa (57.7%) |
| (B) LII - LIV | 1914 | 1867 - 1967 | Sub-Saharan Africa (82.5%) |
| (C) LIV Root | 1964 | 1944 - 1978 | Sub-Saharan Africa (83.5%) |
| (D) LIVme | 1983 | 1976 - 1988 | Middle East (95.4%) |
| (E) LIV asia | 1987 | 1980 - 1992 | Asia (99.1%) |
| (F) LIVne | 1992 | 1984 - 2000 | Sub-Saharan Africa (43.6%) or<br>North-East Africa (48.2%) |
| (G) LIVssa | 1980 | 1967-1993 | Sub-Saharan Africa (90.3%) |

TMRCA, time to most common ancestor; HPD, highest posterior density; Most likely origin, most likely region of origin of the node based on ancestral character reconstruction.

Within LIVssa, three sequences from Nigeria and the Democratic Republic of the Congo (DRC) form a cluster clearly separated from the other sequences in the sublineage. Comparing our phylogenomic results from Nigeria samples with results obtained earlier with the same samples using partial N gene sequences (*14*) (Figure S1), we can identify our cluster of 3 sequences as corresponding to the NigA cluster defined in previous studies (*20, 21*), while the rest of the sequences correspond to the cluster named NigB in the same studies and more widely detected in Nigeria in recent years (*14*). All sequences obtained from PPRV samples in West African countries recently affected by LIV (Mali, Ghana, Côte d’Ivoire and Mauritania) belong to this NigB cluster (Figure 2).

The age of nodes as well as ancestral region of origin were computed alongside the phylogenomics analysis. To decrease bias due to incomplete sampling, node origin modelling was run as per potential region of origin instead of country, setting the regions as Asia, Middle East, sub-Saharan Africa and North-East Africa (Table 2). The root of all PPRV lineages (Node A) was dated to 1828 [95% highest posterior density (HPD) interval: 1744–1908], with sub-Saharan Africa identified as the most likely origin (57.7%). The split between lineages II and IV (Node B) was estimated to have occurred around 1914 (95% HPD: 1867–1967), also originating in sub-Saharan Africa (82.5%).

Within lineage IV, the most recent common ancestor (Node C) was dated to 1964 (95% HPD: 1944–1978), again suggesting a sub-Saharan African origin (83.5%). The subsequent diversification of LIV into regional clades occurred during the late 20th century. The Middle Eastern clade (Node D) was estimated to have a time to most recent common ancestor (TMRCA) of 1983 (95% HPD: 1976–1988), while the Asian clade (Node E) would have diverged shortly thereafter, with a TMRCA of 1987 (95% HPD: 1980–1992). The North-East African clade (Node F) was inferred to have arisen around 1992 (95% HPD: 1984–2000), likely also originating from sub-Saharan Africa (43.6%) or North-East Africa (48.2%). The sub-Saharan African clade (Node G) was estimated to have emerged earliest, around 1980 (95% HPD: 1967–1993).

### Phylogeographical history of LIV in Africa based on partial N gene sequences

The existing dataset of the publicly available PPRV sequence is largest for a small segment of the PPRV nucleoprotein (N) gene used commonly for PPRV diagnosis (*19*). Based on the strong phylogeny produced with full PPRV genomes, this dataset was used to retrace the history of the detection of LIV sequences in Africa, serving as a proxy to provide a first, although highly biased, picture of the spread of this lineage across the continent. The curated dataset of unique N gene sequences available at https://www.ppr-labs-oie-network.org/ was used to perform a Maximum Likelihood phylogenetic analysis and identify African sequences belonging to the sub-Saharan African sublineage IV or to other LIV sublineages (Fig S1). The results show that LIVssa was first detected in Cameroon and Sudan in 1997 and 2000 respectively, then in the Central African Republic in 2004 (Figure 3). It was not reported outside of these 3 countries before 2010, with a first detection in Nigeria in 2010, then South Sudan and Gabon in 2011, and in the Democratic Republic of Congo and Angola in 2012, followed by Niger in 2013. Detections in West Africa continued with Mali in 2017, Senegal in 2020, Burkina Faso in 2021 and, finally, Côte d’Ivoire, Ghana and Mauritania in 2022 (Figure 3). It is worth noting that Sudan is the only country where sequences belonging to both LIVssa and LIVne have been detected, although detection of LIVssa ended in 2008, with LIVne detected only more recently 2016-2017 (Figure 3).

**Figure 3.**
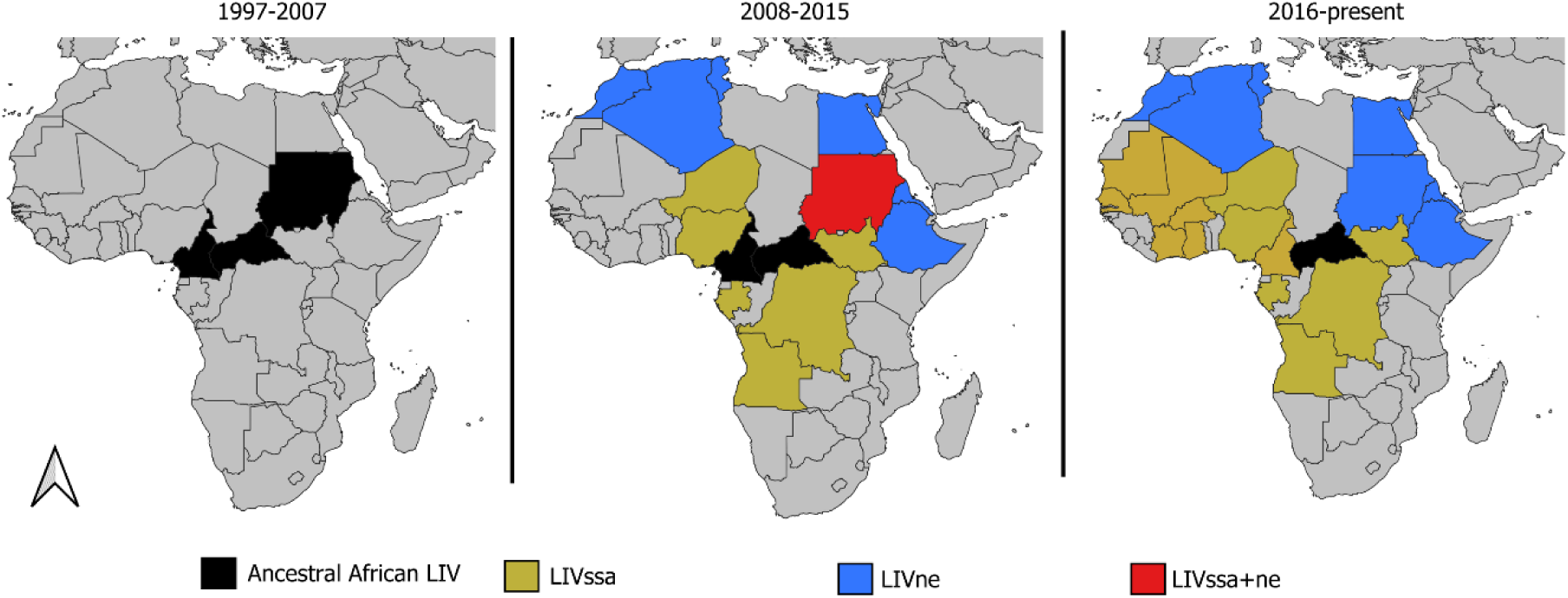
Maps of Africa showing the countries where PPRV genetic lineage IV has been detected at different periods of time between 1997 and 2024. This map is based on publicly available partial PPRV N gene sequences which were identified as belonging to either an ancestral sub-Saharan African or LIV North-East African sublineage IV based on a phylogenetic analysis (see Figure S1). Countries are coloured based on the earliest detection (or confirmed presence) of one or more sequences belonging to these clades within the indicated period. Countries where early sequences not belonging to any clade (basal sequences) have been reported are represented in black.

### Amino acid polymorphisms and selection pressures in the LIV Sub-Saharan Africa group

The recent spread of lineage IV in West Africa could be related to the appearance of genetic mutations modifying the functionality of some viral proteins. This possibility was first explored by comparing the coding sequence of each PPRV gene present in the genomes of strains belonging to the lineage IV sub-Saharan Africa clade and other sequences available. Multiple codon positions in all genes except the M gene coded for amino acids observed only in the lineage IV sub-Saharan Africa group, with the highest number of polymorphisms (n=8) identified in each of the H and L genes (Table 3; Figure 4). A total of 22 amino acid changes specific to the recent (since 2000) sequences of this group could also be identified across all genes (Table 3; Figure 4). Interestingly, two polymorphisms were observed in sequences of the clade NigB only, with one change from methionine to threonine at position 472 of the nucleoprotein, and one change from phenylalanine to leucine at position 220 of the haemagglutinin, also detected in two sequences from Israel (Table 3; Figure 4). Almost half of these codon changes (22 out of 48) were associated with a change in charge, hydrophobicity or size of the amino acid, potentially affecting their functionality (Table 3).

**Figure 4.**
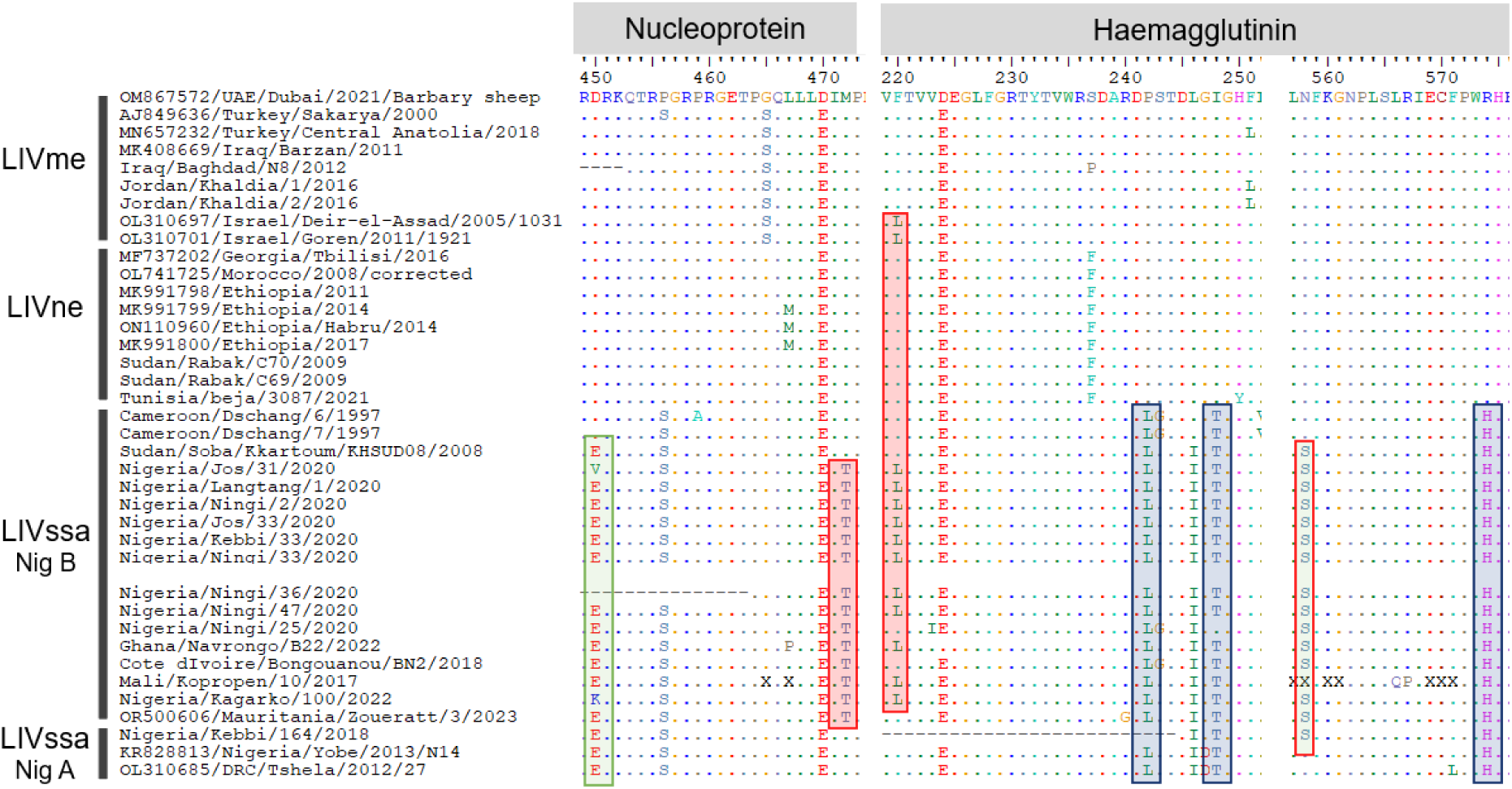
Partial alignment of amino acid sequences of the nucleoprotein (N) and haemagglutinin (H) of peste des petits ruminants virus (PPRV), highlighting examples of amino acid polymorphisms present in all sequences within the lineage IV sub-Saharan Africa group (LIVssa; blue box), in recent sequences (>2000) of this group (green box) and in the clade NigB of the LIVssa group only (red box). Sequences presented in this figure are identified as belonging to different lineage IV geographic groups based on the phylogeny presented in Figure 2.

**Table 3.**
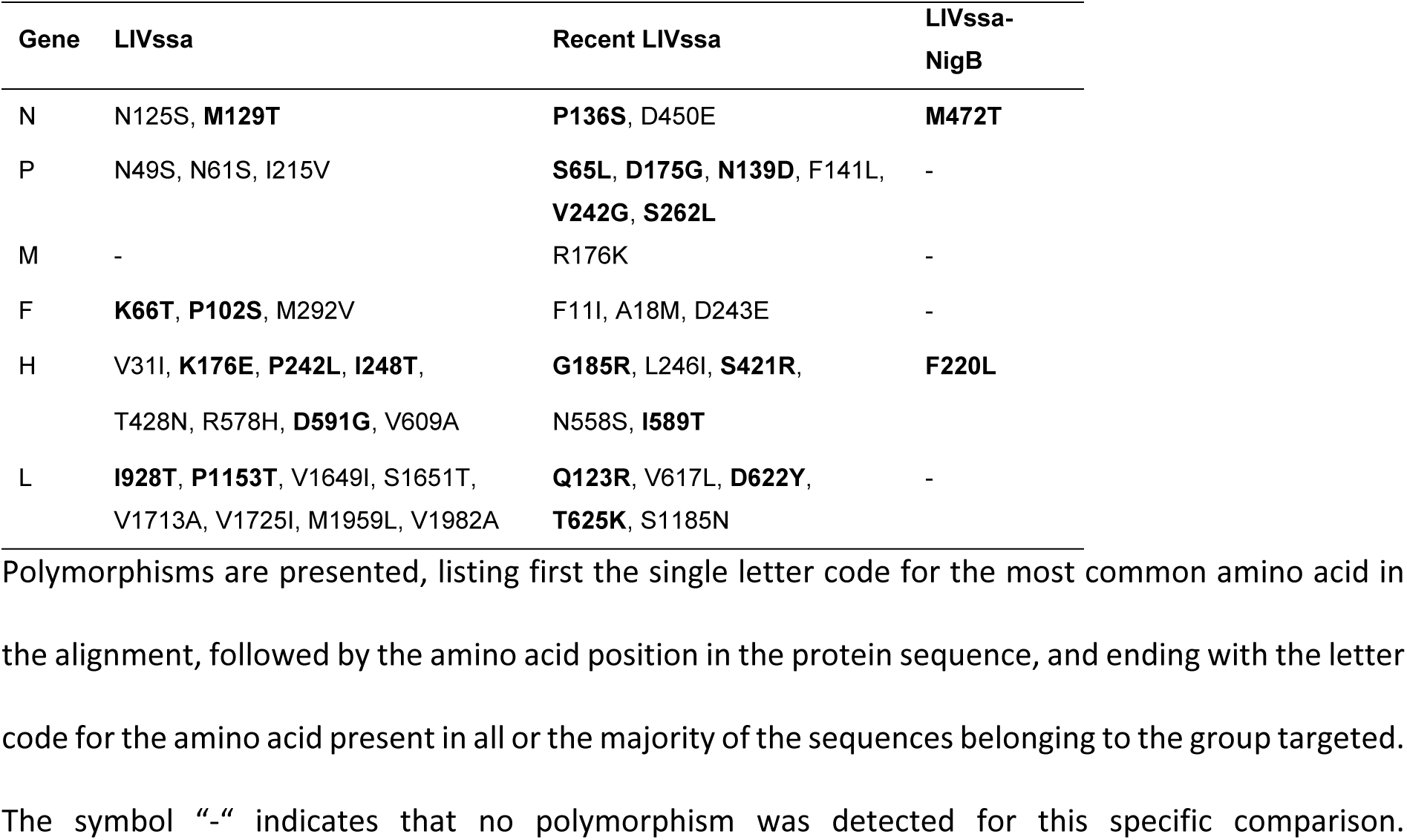
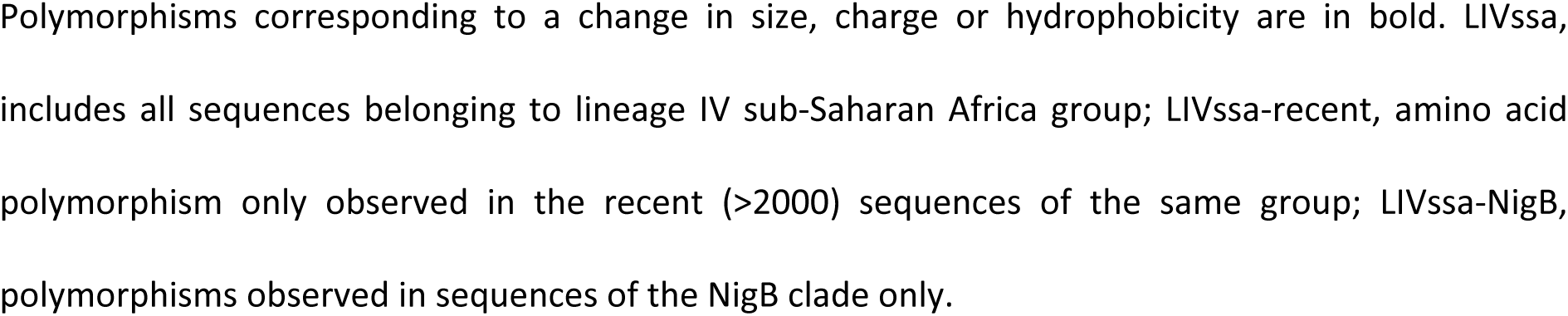
Amino acid polymorphism between sequences of lineage IV sub-Saharan Africa group (LIVssa) and other lineage IV groups.

| Gene | LIVssa | Recent LIVssa | LIVssa-NigB |
| --- | --- | --- | --- |
| N | N125S, **M129T** | **P136S**, D450E | **M472T** |
| P | N49S, N61S, I215V | **S65L**, **D175G**, **N139D**, F141L, **V242G**, **S262L** | - |
| M | - | R176K | - |
| F | **K66T**, **P102S**, M292V | F11I, A18M, D243E | - |
| H | V31I, **K176E**, **P242L**, **I248T**,<br>T428N, R578H, **D591G**, V609A | **G185R**, L246I, **S421R**,<br>N558S, **I589T** | **F220L** |
| L | **I928T**, **P1153T**, V1649I, S1651T,<br>V1713A, V1725I, M1959L, V1982A | **Q123R**, V617L, **D622Y**,<br>**T625K**, S1185N | - |

Polymorphisms are presented, listing first the single letter code for the most common amino acid in the alignment, followed by the amino acid position in the protein sequence, and ending with the letter code for the amino acid present in all or the majority of the sequences belonging to the group targeted. The symbol “-” indicates that no polymorphism was detected for this specific comparison.
Polymorphisms corresponding to a change in size, charge or hydrophobicity are in bold. LIVssa, includes all sequences belonging to lineage IV sub-Saharan Africa group; LIVssa-recent, amino acid polymorphism only observed in the recent (>2000) sequences of the same group; LIVssa-NigB, polymorphisms observed in sequences of the NigB clade only.

Secondly, selection pressure in the sub-Saharan African LIV clade was explored in the coding region of each PPRV gene using multiple methods. BUSTED (branch-site unrestricted statistical test for episodic diversification) was used to identify gene-wide evidence of positive selection in the LIV sub-Saharan Africa group. Significant evidence of episodic diversifying selection was detected only for the N gene in this group (LRT=19, *p*-value < 0.001). The RELAX method was used to test for the difference in selection pressure between LIVssa group and other LIV geographic groups. A difference in selection pressure was identified for the L gene (LRT=11.59, p-value = 0.001) using this method. Results for both RELAX and BUSTED were not significant for the other PPRV coding regions. Fixed effect likelihood (FEL) analysis was run for the coding region of each gene to detect site-specific selection pressure in the LIVssa group, focusing on either all recent sequences (>2000) or only the sequences of the NigB. Multiple codon positions were identified as being under diversifying selection in all genes of the NigB clade and in nearly all genes (excluding M and F) across the recent sequences of the LIVssa group (Table 4).

**Table 4.**
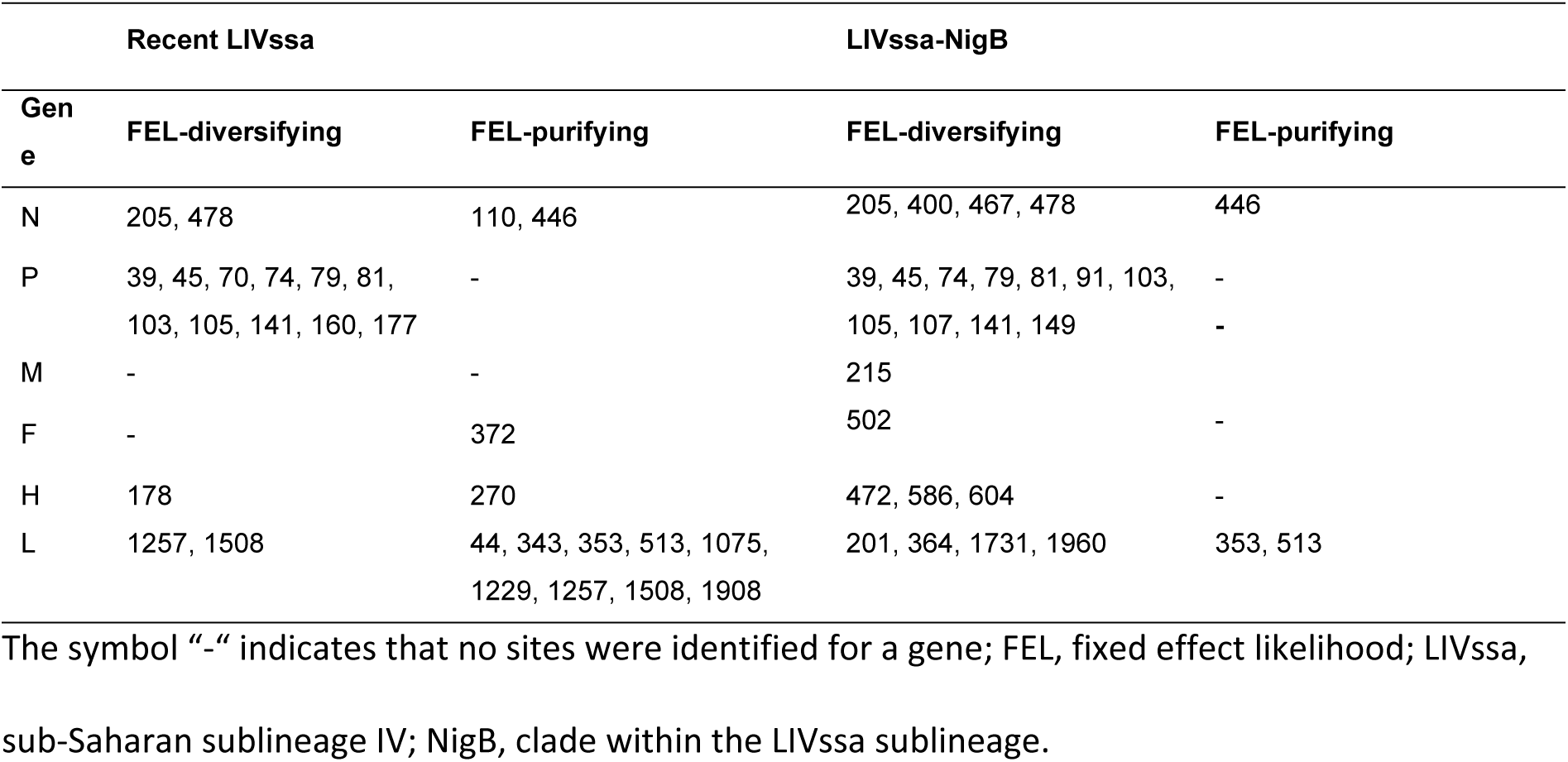
Codon positions under diversifying or purifying selection pressure in the lineage IV sub-Saharan Africa group.

| Gene | Recent LIVssa |  | LIVssa-NigB |  |
| --- | --- | --- | --- | --- |
|  | FEL-diversifying | FEL-purifying | FEL-diversifying | FEL-purifying |
| N | 205, 478 | 110, 446 | 205, 400, 467, 478 | 446 |
| P | 39, 45, 70, 74, 79, 81, 103, 105, 141, 160, 177 | - | 39, 45, 74, 79, 81, 91, 103, 105, 107, 141, 149 | - |
| M | - | - | 215 | - |
| F | - | 372 | 502 | - |
| H | 178 | 270 | 472, 586, 604 | - |
| L | 1257, 1508 | 44, 343, 353, 513, 1075, 1229, 1257, 1508, 1908 | 201, 364, 1731, 1960 | 353, 513 |

The symbol “-” indicates that no sites were identified for a gene; FEL, fixed effect likelihood; LIVssa, sub-Saharan sublineage IV; NigB, clade within the LIVssa sublineage.

Sites under selection were most numerous for the P gene in both analyses. The sites identified were similar for the N and P genes in the two groups but not for H and L genes (Table 4). Evidence of purifying selection was also detected in some positions for all genes except P and M in the recent sequences of the LIVssa group, but only in two codon positions in the N and L genes when the analysis was run solely on the NigB clade (Table 4).

## Discussion

Despite ongoing efforts to eradicate peste des petits ruminants, the risks of the spread of this devastating disease remain extremely high given the current context of global trade and political instability. The recent emergence of PPR in Europe is a clear example of this risk (*3*), but the ongoing spread of the PPRV lineage IV in new regions of Africa also shows that the context of endemic PPR circulation in sub-Saharan Africa may have changed. The fact that PPRV lineage IV is currently extending its distribution in West Africa has been well recorded since its detection in Mali in 2017 (*11–13, 15, 16*), but the earlier history of the presence of lineage IV in Africa is much more confused. The most common story presented is that lineage IV originated in Asia and spread to the Middle East and Africa (*22, 23*), but more recent phylogenomic analyses have suggested an African origin for LIV, despite the paucity of data available (*4, 9*).

Here, with 26 new full or almost complete PPRV LIV genomes from Africa, our phylogenomic analyses provide strong support for an African origin of lineage IV. Several, geographically separated, sublineages within LIV had been identified in the past (*4*), and here we are able to add a new sublineage corresponding to strains circulating in sub-Saharan Africa (called here LIVssa). Our analyses show that this cluster is the most ancestral LIV group. Based on publicly available partial N gene data, we have confirmation that lineage IV has been present since at least 1997 in Cameroon, and it spread from this region to other parts of Africa. However, sequence data is too limited to provide a strong narrative for the early distribution of lineage IV in Africa. It was certainly present in the region before 1990 since lineage IV has been recorded since at least 1987 in India (*24*). The estimation of the TMRCA for LIV suggests the lineage appeared at the latest in 1978, with a divergence between LII and LIV starting at the latest in 1967 (highest 95% HPD value, see Table 2). These estimations differ by only 2-10 years to those obtained in earlier studies with smaller genome datasets and are accompanied large uncertainties (*4, 9*), but all fit with an appearance of LIV in Africa before 1990. Obtaining genomic sequences from additional historical samples, as we have done with Mali (1999) and Cameroon (1997) may refine these estimations, but there is little probability that new sequence data would put the African origin of LIV into question.

Data from Sudan provide an interesting insight into the evolutionary history of lineage IV in Africa, as it is the only country where two different LIV sublineages (LIVssa and LIVne) have been detected. Based on partial N gene sequences, we know that LIVssa has been circulating since at least 2000, at the same time as LIII, and that both LIVssa and LIVne were circulating in 2008. Only LIVne sequences have been published since then in Sudan, suggesting that LIVssa strains have had no competitive advantage over LIVne strains. This LIVne sublineage appears to be more recent than the Middle East sublineage LIV, suggesting a movement of ancestral LIV strains into the Middle East and then to Asia, before the introduction of new LIV strains from Asia and the Middle East to African countries bordering the Red Sea and the rest of North Africa, leading to sublineage LIVne. Here again, more genetic data from these poorly sampled regions would help to investigate this hypothesis further. Additionally, it would be interesting to assess if historical patterns of animal legal and illegal movements across these regions fit with the genetic results. Lineage III seems to have been replaced by LIV in Sudan but no such trends have been observed in East and Central African countries where LIII is circulating endemically (*17*), although this possibility may need to be investigated more thoroughly. In this study, we were able to obtain two additional LIII genomes from Sudan and Comoros, but this lineage and lineage I remain understudied compared to LII and LIV.

Nigeria is the only country where there is enough genetic data to suggest a dominance of LIV over LII in countries where strains of both lineages are circulating (*14, 25*). Distribution of both lineages in other West Africa countries where LIV has recently emerged will need to be carefully followed to assess whether the LIV strains are replacing LI and LII strains where they typically circulate endemically. The ongoing detection of LII in Nigeria and LI in Mali suggests that, even if a new lineage may come to dominate in a region, lineages historically present may remain in some areas and only thorough sampling would allow confirmation of the disappearance of a lineage in a region.

Genetic data suggests that the NigB group within LIVssa is more widely distributed than the NigA group (*14*). The latter group has been detected in neighbouring African countries (Gabon, RDC, Niger and Angola), with the latest reports in Niger in 2015 and Nigeria in 2018. However, our phylogenomic results clearly show that PPRV strains associated with the recent spread of LIV in West Africa belong to the NigB cluster, suggesting this group may have acquired some mutations that have facilitated its spread and dominance over LII and LIV NigA strains. We have found substantial evidence of selection pressure on the LIVssa sublineage and, more specifically, on the NigB cluster in all PPRV proteins. Some amino acid changes specific to LIVssa have also been identified in all PPRV proteins, including two changes specific to NigB in the nucleoprotein and haemagglutinin. Although none of these codon positions are directly linked to a specific role in interactions with other PPRV or host proteins, they may lead to changes in protein function. Most codons under selection or with amino acid changes specific to LIVssa were observed in the N terminal region of the phosphoprotein, responsible for binding to the nucleoprotein (*26*), in the putative functional site of the RNA-dependent RNA polymerase code (*27*), and in the haemagglutinin, although mostly in sites distant from the region interacting with the host cellular receptor (aa505-552) (*28*). Based on the hypothesis that the NigB group has evolved some adaptive advantages leading to its spread in West Africa, the impact on protein function of the two mutations detected in the nucleoprotein (methionine to threonine at position 472) and haemaggluttinin (phenylalanine to leucine at position 222) should be investigated as a priority.

However, further genomic data is needed to confirm the association between the NigB group and the current spread of LIV in West Africa. This spread may have been facilitated by other mutations differentiating the recent strains within the LIVssa clade from other LIV strains. Other socio-economic factors may also have facilitated the spread of lineage IV in the past decade, with several issues such as droughts or conflicts that may change trade patterns or drive the transboundary movement of people and their animals. Integrative studies are therefore needed to fully understand the current changes in PPR circulation dynamics.

This study only starts to fill the gaps in our understanding of the evolutionary history of the most widely distributed genetic lineage of PPRV. Our results clearly show that sub-Saharan Africa has consistently been the hotbed for PPRV evolution, including for lineage IV. More genomic work needs to focus on historical and recent viral strains circulating in this region to provide some insights that may help current efforts to eradicate the disease. Genomic data confirms that the spread of LIVssa in West Africa is a concern and should be the centre of more research efforts. Indeed, if PPRV strains within LIVssa have evolved in a way that facilitates their transmission through faster replication, capacity to infect a wider diversity of hosts or other modifications that increase their fitness, this may affect the current effort to control the disease in regions where its impact on the economy and food security is the highest. The infection capacity of LIVssa in sheep and goats needs to be urgently compared to other strains circulating endemically in sub-Saharan Africa in experimental settings to evaluate the need to review control strategies, notably the minimum vaccination coverage to be targeted in order to stop virus circulation.

## Material and Methods

### Sample preparation and high-throughput sequencing analysis

Samples received and treated at the Joint FAO/IAEA Centre were prepared following a method described in detail elsewhere (*12*), based on the preparation of the sequencing library by sequence independent single primer amplification (SISPA) from total nucleic acid extracted from homogenised samples using the RNeasy Mini Kit (Qiagen, Germany), before sequencing on a S5^TM^ next-generation sequencing system (Ion Torrent, Thermo Fisher). At CIRAD, multiple methods of sample preparation have been implemented across the years of genome sequencing efforts (Table 1). Several samples were prepared and analysed by high-throughput sequencing using either a modified SISPA protocol (*29*) or a protocol based on the PCR amplification of PPRV fragments (*30*) followed by HiSeq or MiSeq Illumina sequencing as described elsewhere (*4*). The majority of samples processed at CIRAD for this study were first treated with a mix of nucleases following a protocol described in (*31*) before extraction of nucleic acids using a magnetic beads-based extraction kit (IndiMag Pathogen INDICAL BIOSCIENCE) according to the manufacturer’s instructions, except for the replacement of the carrier by 1µl of linear acrylamide (5 mg/ml, Invitrogen). For the preparation of the library, first strand cDNA was generated with random hexamer primers using the RevertAid first strand cDNA synthesis kit (Thermo Scientific) according to the manufacturer’s protocol, except for the addition of 1µl of 100% DMSO at the denaturation step. After a step of digestion at 37°C for 20 minutes with 2U of RNase H, double stranded cDNA was synthesised using the DNA Polymerase I, Large (Klenow) Fragment kit (New England Biolabs) in a final volume of 30μl incubated at 25 °C for 15 minutes followed by 75° C for 20 minutes. Double stranded cDNA was purified using the Nucleospin PCR clean up and gel extraction kit (Macherey-Nagel) according to the manufacturer’s instructions in a final volume of 30µl. The Qubit dsDNA HS assay kit (Invitrogen) was used to evaluate the concentration of the purified products and adjust the concentration to 0.2 ng/μl. Libraries were generated using the Nextera XT DNA Library Prep kit according to the manufacturer’s instructions in ‘dual index’ by using two indexes for each sample. Concentration of the libraries was assessed with the library quantification kit (Takara Clonetech) and distribution of fragment sizes analysed on an Agilent 2100 bioanalyser with the Agilent high sensitivity DNA kit (Agilent Technologies). Based on the results obtained, libraries were pooled in an equimolar manner before performing sequencing on a MiniSeq Illumina machine at the AGAP or MGX sequencing platforms (Montpellier, France).

### Sequence data analyses and phylogenetic analyses

The raw sequences obtained were cleaned and processed to generate a consensus genome sequence following a bioinformatic pipeline described in (*12*) for the Joint FAO/IAEA Centre, and in (*4*) for CIRAD. The absence of contamination in all consensus sequences generated was confirmed using the multiple recombination tests implemented in RDP5 (*32*) as described in (*19*). We used TempEst v1.5.3 (*33*) to verify that our data evolved in a clock-like manner and to detect any outlier sequence that did not fit with the overall association between genetic divergence and sampling date.

Phylogenetic analyses were performed using an uncorrelated relaxed molecular clock model with a lognormal distribution in combination with a coalescent constant population size prior (*34*). Nucleotide substitution was modelled under the general time reversible (GTR) model with four gamma-distributed rate categories. The ancestral regions of origin for the internal nodes were estimated with a discrete trait reconstruction alongside the tree estimation during the Bayesian analysis in Beast. Sampling locations for tips, based on both extant and historical reports of PPR occurrence, were categorised into four geographic regions: Asia, Middle East, sub-Saharan Africa and North-East Africa. We used a discrete trait substitution model with asymmetric transition rates among all regions, implemented with Bayesian Stochastic Search Variable Selection. Two independent Markov Chain Monte Carlo (MCMC) runs of 500 million steps each were performed. Convergence and effective sample sizes (ESS) of parameters were assessed in Tracer v1.7.1 (*32*). Posterior distributions of node ages, evolutionary rates and region of origin probabilities were summarised by extracting quantiles from the log files in R (*33*).

### Amino acid polymorphism and detection of selection pressures in sub-Saharan African lineage IV clade

Alignments were created for the coding region and amino acid sequences of each PPRV gene. Amino acid alignments were manually searched for the presence of polymorphisms unique to the sequences belonging to lineage IV sub-Saharan Africa group. Polymorphism present in all but one or two LIVssa sequences or present also in a few sequences belonging to another group were noted. In the same manner, specific amino acid changes only observed in the recent (>2000) sequences of LIVssa and in the LIVssa-NigB were listed separately.

Three different methods were implemented through the datamonkey.org platform (*34*) to assess selection pressure in the coding sequences of each PPRV gene, focusing on the sequences of the lineage IV sub-Saharan Africa group. First, gene-wide evidence of episodic selection on the branches of LIVssa was evaluated with the BUSTED (branch-site unrestricted statistical test for episodic diversification) approach (*35*). A second method called RELAX (*36*) was used to test if the strength of selective pressure differed between LIVssa and other LIV groups. Finally, a maximum-likelihood approach called fixed effect likelihood (FEL) (*37*) was implemented for each gene to detect individual codon positions under diversifying or purifying selection, based on nonsynonymous (dN) and synonymous (dS) substitution rates, and assuming that the selection pressure on each site is constant along the phylogeny. FEL was run on two sequence subsets most likely associated with the recent spread of LIV in West Africa: all recent (>2000) sequences of LIVssa and sequences of the NigB clade.

## Acknowledgements

AB is supported by a grant from the European Commission Animal Health and Welfare European Research Area Network for the IUEPPR Project ‘Improved Understanding of Epidemiology of PPR’, in the framework of ANIHWA 2013, and by a grant (SI2.756606) from the European Commission Directorate General for Health and Food Safety awarded to the European Union Reference Laboratory for peste des petits ruminants (EURL-PPR). Part of this work was supported by the VETLAB network initiative of the FAO/IAEA Centre.

## Data availability

Raw genetic data was deposited in the NCBI Sequence Read Archive (accession number: PRJNA717034), and consensus genomes sequences were deposited in NCBI GenBank (accession numbers: PX459241, PX460960- PX460962, PX685403-PX685424).

## Supporting information

**Figure S1.** Consensus phylogenetic tree of partial PPRV nucleoprotein gene sequences obtained using Maximum Likelihood method. Numbers at tree nodes correspond to the percentage of confidence support based on bootstrap analysis (1,000 replications). Only nodes with >50% support are presented. Sequences from major genetic lineages and sublineages with no direct relevance for this analysis have been collapsed so only LIV sequences from African samples are shown. The complete tree is available in Newick format.

**Figure S2.** Correlation between root-to-tip genetic divergence and sampling date in the PPRV genome dataset as calculated using TempEst.

